# Relative electrical conductivity measurements of the rat frontal cortex and the influence on current source density

**DOI:** 10.1101/128322

**Authors:** Yevgenij Yanovsky, Jurij Brankačk

## Abstract

The relative electrical conductivity gradient with depth was estimated in the frontal cortex of anaesthetized rats. Current source density (CSD) approximations of field potentials evoked by ventromedial thalamic stimulations with an assumed homogeneous electrical conductivity of the neocortical tissue were compared to those with correction for the estimated conductivity gradient. In spite of the cellular heterogeneity the electrical conductivity of the frontal cortical tissue was found to be fairly homogeneous inside the superficial (layers I through IV) or deep layers (V- VI). The relative conductivity increased twofold at the transition between superficial and deep layers. Regardless of this changes CSD analysis of the field potentials evoked by ventromedial thalamic stimulation revealed negligible differences between estimations ignoring the conductivity and those taking the conductivity into account. No sinks or sources appeared or disappeared. Both CSD approximations revealed: 1) a strong sink in layer I representing most likely summed monosynaptic EPSPs of the ventromedial thalamic afferents; 2) a strong sink in layer VI, probably representing summed disynaptic EPSPs on dendrites of layer VI pyramidal cells, generated by axons of upper layer pyramidal cells; and 3) a sink in lower layer V representing probably threesynaptic summed EPSPs on dendrites of layer V pyramidal cells.

## 1. Introduction

Current source density (CSD) analysis is a powerful method for location of transmembrane currents from a set of extracellular field potentials recorded at different depths [6,14,17]. In combination with additional data it allows to identify active synaptic sites. The CSD method was successfully applied in a number of laminated brain structures mainly for studies on evoked activity [13], seizure activities [2,16] and EEG rhythms [3,5,16,17].

Large conductivity variations of the cortical tissue with depth can lead to increased noise levels of the CSD and possibly produce spurious sinks and sources [14,17]. Therefore it is desirable to know the degree of conductivity changes with depth and their influence on CSD estimations. Rappelsberger and colleagues [17] working on rabbit’s visual cortex have emphasized the need of conductivity measurements for CSD analysis in every experiment whereas others have denied significant influences on CSD calculations [7,12,19]. Holsheimer [11] investigated the conductivity of hippocampal CA1 layers in guinea pigs and found a drop of the electrical conductivity to as much as 42% in the pyramidal cell layer compared to the stratum radiatum, a mainly dendritic layer. In spite of this differences in conductivity the spatial distribution of sinks and sources did not differ between CSD approximations with homogeneous and inhomogeneous conductivity [11]. The neocortical tissue seems to be even more homogeneous compared to the hippocampal CA1 area where the pyramidal cell bodies are densely packed in a single layer. The conductivity gradient with depth and its influence on CSD analysis could be possibly weeker in the neocortex than in the hippocampus. In addition, there are cytoarchitectonic differences between layers of the primary visual cortex and those of the frontal cortex. Layer IV in the primary visual cortex is large and more distinct than in the frontal cortex where it is indistinguishable from layers II-III [18]. The bodies of layers V and VI pyramidal cells of the frontal (motor) cortex are larger in size compared to those of the visual cortex. No data is available about electrical conductivity of the rat frontal cortex neither about current source density analysis. The present experiments were designed to study the degree of relative conductivity changes with depth in the rat frontal cortex and its significance on CSD estimations. Portions of this work have been published in abstract form [4].

## 2. Materials and methods

### 2.1. Animals and surgery

Fifteen 4-6 month old male Wistar rats (270-530g) were anaesthetized with urethane (i.p., 1.2 g/kg) or a mixture of pentobarbital (i.p., 40 mg/kg) and urethane (lOmg/kg) and mounted in a stereotaxic instrument with bregma and lambda at the same height. A pair of 100 μm diameter varnish insulated nichrome wires (cut square with 1.0 mm vertical tip separation) was implanted in the ventromedial thalamic nucleus (VM) to stimulate thalamo-cortical fibers (2.3 mm posterior from bregma, 6 mm ventral to bregma, 1.8 mm lateral from midline [15]), contralateral to the cortical recording electrode. An approximately 3.5 mm diameter hole was drilled in the skull above the frontal cortex, area Frl [18] with its center from 0.0 mm to 2.5 mm anterior to bregma and 2.0 mm to 3.0 mm lateral to the midline. The dura mater was removed at the recording site. A nylon tube (5.0 mm inner diameter) was fixed above the hole with dental acrylic and filled with agar gel (see 1 in fig. 1).

### 2.2. Conductivity measurements

The relative electrical conductivity was estimated at multiple points throughout the depth of frontal cortex (area Frl) using tungsten microelectrodes (3 MΩ) moved slowly in steps of 50 μm. A sine wave (1,000 Hz; 0.3 mV) was applied monopolarly to a ring-shaped silver wire (4.00 mm ring diameter, 0.4 mm wire diameter) which was placed into the agar filled nylon tube (1, fig. 1) and mounted on the cortical surface, so that the silver electrode was situated 6 mm above the cortical surface (2). The second electrode was placed into the rat’s mouth and pressed toward the upper palate (3). It consisted of a stainless steel plate (70 square mm) wrapped in saline soaked filter paper. The sine wave signal had no effect on frontal cortical activity. The voltage recorded at each microelectrode step in reference to a distant indifferent electrode at the temporal bone of the rat’s scull (4) was fed to a 1kHz pass filter. Averaged voltage measurements (N=10) at each step were plotted as a function of depth. With current constant, the difference in voltage recorded at two neighbouring points, i.e. the slope of the voltage decrease with depth, is proportional to the tissue conductivity. A change in the slope stands for a change in conductivity. For averages among animals the voltage potentials were normalized to the potential at the cortical depth of 750 μm, approximately the middle of the cortex [20], which was set to 1.0.

**Fig. 1:**
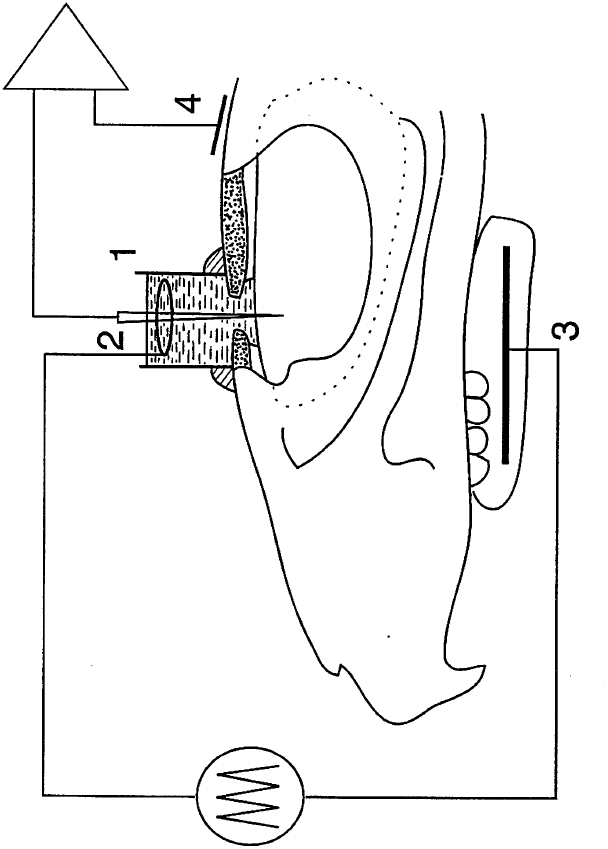
Scheme of the technique for in vivo measurements of the relative cortical tissue conductivity. ‘1’: nylon tube filled with agar gel; ‘2’: silver ring electrode; ‘3’: stainless steel plate in the rat’s mouth, which was wrapped in saline soaked filter paper and pressed toward the upper palate; ‘4’: reference electrode. A 1 kHz sine wave (0.3 mV) was applied to electrodes ‘2’ and ‘3’. The signal between the microelectrode and the reference electrode was recorded at 50 μm steps and fed to a 1 kHz pass filter.

### 2.3. Current source density analysis

At each step of the microelectrode, evoked field potentials to ventromedial thalamic stimulations (0.1 ms pulse duration; from 0.5 mA to 1.5 mA, presented once every two seconds) were recorded, filtered (1 Hz high pass and 2 kHz low pass filter), digitized at 10 KHz, and averaged. The data were stored on disk for off-line analysis. Voltage profiles as a function of recording depth were computed from the averaged evoked potentials by smoothing them three times to reduce the high spatial frequency noise. Such smoothing was performed with a simple three point rolling average of the voltage in depth using a weighting factor of two for the center point and one for each of the adjacent points [3,6]. The second differences of the voltage profile were divided by the square of the step size in cm. The current source density (CSD) analysis was performed in one dimension (depth) for field potentials evoked by stimulation of the ventromedial thalamic nucleus. The measured relative cortical tissue conductance for each step was used for computing the conductivity corrected approximations of current source densities. One-dimensional CSD analysis was performed according to the procedure described by Nicholson and Freeman [14]:

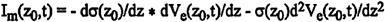

where I_m_(z_0_,t) is the approximation of the transmembrane current density at the location z_0_ and the time t, σ(z_0_) represents the electrical conductivity of the tissue at the location z_0_, and V_e_(z_0_,t) the field potential at the location z_0_ and the time t. a) If the tissue conductivity is considered to be homogeneous and the field potentials V_e_ are recorded at discrete spatial intervals h the current density is estimated by the following equation:

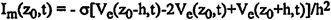

b) For an inhomogeneous tissue conductivity

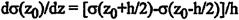

the equation will be the following [11,17]:

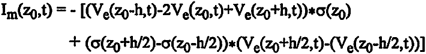

In order to better visualize current source density (CSD) as a function of both time and depth within the profile, contour maps were plotted. Smooth contours were generated by the computer, using linear interpolation between data points. Contours representing sinks were coded with solid lines and those representing sources with dashed lines.

## 3. Results

The electrical conductivity gradient with depth was estimated in the rat frontal cortex by using the technique described in the methods section (see fig. 1). A change of the voltage potential slope with depth indicates changes of the tissue conductivity. Fig. 2 shows the normalized potential (ordinate) as a function of depth (abscissa) averaged among six animals.

**Fig. 2:**
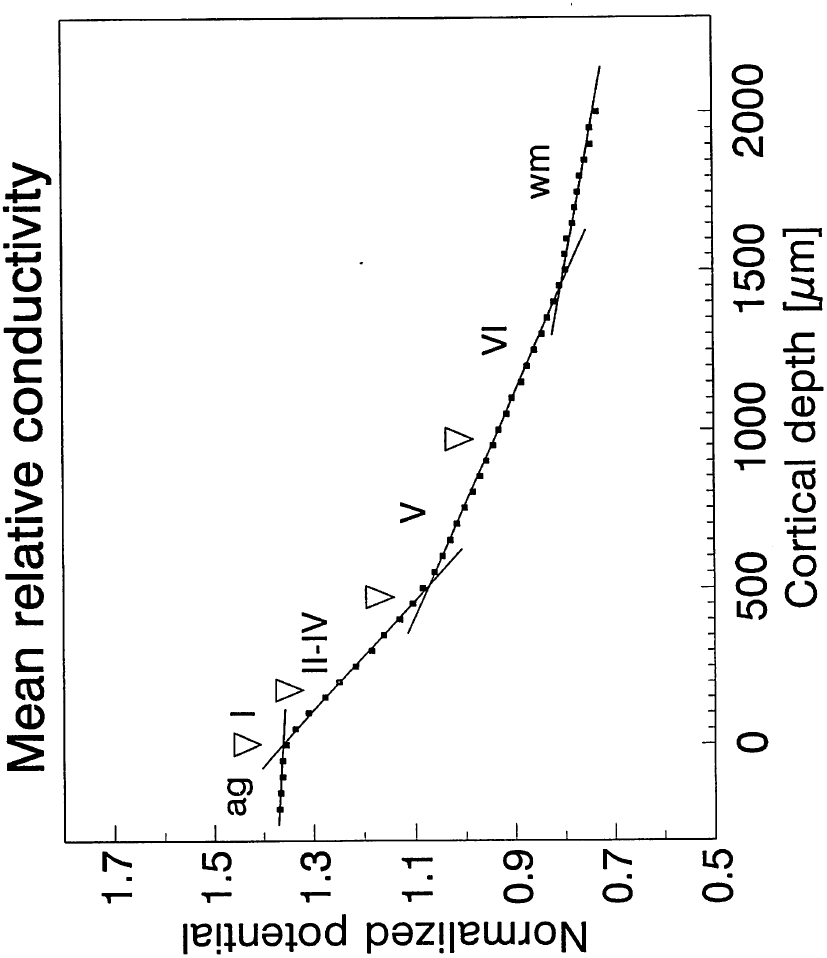
Conductivity related potentials, normalized to a depth of 750 μm (=1.0) and averaged among animals (ordinate) are shown as a function of depth (abscissa). White triangles mark the cortical surface and boundaries between frontal cortical layers I through VI; ‘ag’: agar gel; ‘WM’ white matter.

White triangles mark boundaries between the cortical layers [18]. Four regression lines are drawn: 1) line I (−200 to 0 μm, corresponding to agar gel (ag) above the cortical surface): r= −0.949, p<0.01; 2) line II (0 to 450 μm, corresponding to the superficial layers I-IV): r= −0.999, p<0.01; 3) line III (500 to 1400 μm, corresponding to the deep layers V and VI): r = −0.999, p<0.01; and 4) line IV (1450 to 2000 μm, corresponding to white matter (wm)): r= −0.990, p<0.01. The least conductivity (steepest slope of the potential change with depth) was found for the superficial layers followed by the deep layers which had a twofold higher conductivity than the superficial layers (49.9% of deep layers) and the white matter with an about twofold higher conductivity than the deep layers of the frontal cortex (45.8% of white matter). The conductivity of the agar gel was nine times larger than in the superficial cortical layers (11.1% of agar gel). Inside the frontal cortex conductivity changes are seen on the transitions between superficial layers I-IV and the deep layers V-VI and between the deep layer VI and the underlying white matter.

Fig. 3 shows the influence of conductivity changes with depth on current source density estimations in a representative electrode track. Part A displayes the depth profile of field potentials evoked by stimulation of the ventromedial thalamic nucleus. Figures on the left indicate cortical depth (in μm). The arrow marks the thalamic stimulation. In part B the corresponding conductivity related potential (ordinate) change with cortical depth (abscissa) is shown. Parts С and D show contour plots of current source density (CSD) estimations of the field potentials (shown in part A). Cortical depth is represented by the ordinates. The abscissae correspond to the time after thalamic stimulation (0 ms). Solid lines indicate sinks and dashed lines sources. In part C a homogeneous conductivity was asumed and in D the conductivity related potential gradient in B was used for correction. The distance between contour lines for both sinks and sources in part C corresponds to 3 mV/mm^2^ and in D to 15.0 mA/mm^3^. Although there are minor differences between the two contour plots, no sinks or sources appear or disappear in C compared to D. The CSD in C and D shows: 1) week inward membrane currents (sink) at about 0.3 ms in layer VI which represents most likely antidromic activation of the cortico-thalamic axons of layer VI pyramidal cells; 2) a strong sink with a latency of about 0.5 ms in layer I (slightly displaced towards layer II) drowing its current from outward membrane currents (sources) above and below: Most likely it represents the summed monosynaptic EPSPs of ventromedial thalamic afferents projecting to layer I. The depth shift (50 to 100 μm) of the sink could be due to mechanical displacement of the cortical surface by the electrode entering the cortex, 3) a strong sink with a latency of 1.5 ms in layer VI coupled with sources above and below corresponds to summed EPSPs of interlaminar projections of layer I/II pyramidal cells to layer VI pyramidal cells, 4) a third strong sink with a latency from 2 to 2.5 ms in layer V/layer VI coupled with a source in layer V represents probably summed EPSPs on layer V/VI pyramidal cells.

**Fig. 3:**
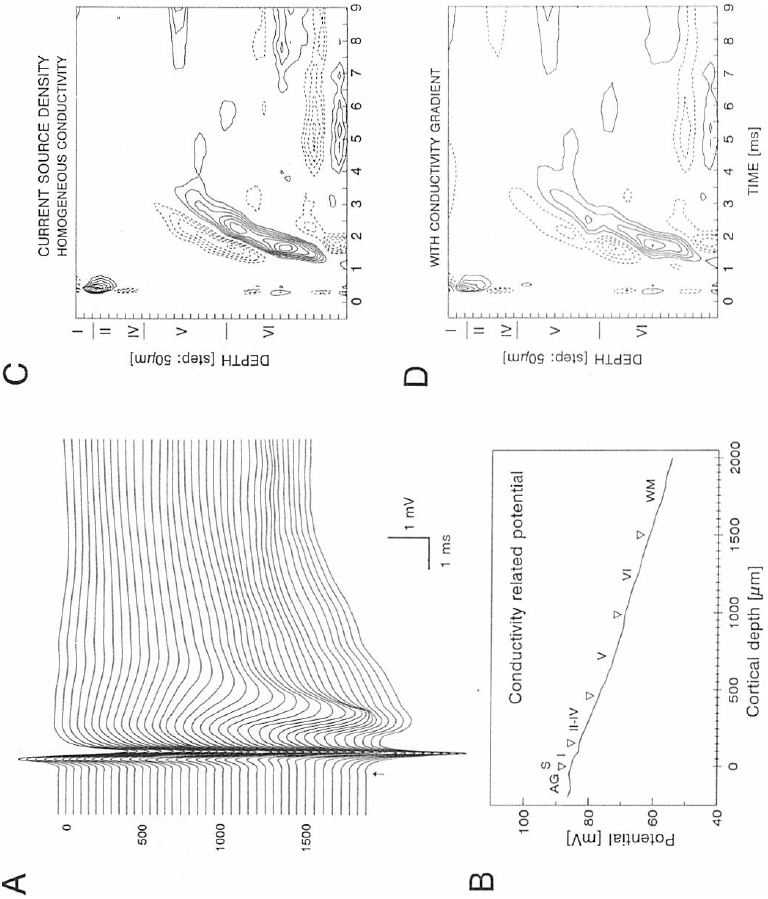
Influences of electrical conductivity changes with depth on current source density estimations shown on a representative electrode track. **Part A:** Depth profile of field potentials in the frontal cortex evoked by stimulation (arrow) of the ventromedial thalamic nucleus. Figures on the left indicate the cortical depth in μm. Positive is up. **Part B:** The conductivity related potential measured on each electrode step (for details see methods and results section). The triangles show the boundaries between agar gel (AG) and the cortical surface (S), between the cortical layers (I to VI) and between layer VI and the underlaying white matter (WM). The cortical layers were estimated for an average frontal cortical thickness of 1.5 mm: layer I: 163 μm (11%), layer II-IV: 300 μm (20%), layer V: 514 μm (34%) and layer VI: 523 mm (35%) [18,20]. **Part C and D:** Contour plots of current source density (CSD) estimations of the field potential profile in A. Solid lines represent sinks and dashed lines indicate sources. **Part C** displayes the CSD estimations ignoring changes of the electrical conductivity. Amplitudes in C: for sinks from ‒6.0 to ‒27.0 mV/mm^2^, and for sources: from 6.0 to 18 mV/mm^2^. The distance between contour lines for both sinks and sources: 3 mV/mm^2^. **Part D** the CSD was corrected for conductivity changes with depth using potential differences between steps shown in part B. Amplitudes in D: for sinks from ‒15.0 to ‒90.0 mA/mm^3^ and for sources from 15.0 to 90.0 mA/mm^3^, The distance between contour lines for both sinks and sources: 15.0 mA/mm^3^.

## 4. Discussion

The main findings of the present study are, 1) conductivity changes with depth in the frontal cortex are restricted to the boundary between superficial and deep cortical layers: the average conductivity of the deep layers is two times higher, 2) in spite of a twofold increase of conductivity there are negligible differences between current source density (CSD) estimations including the conductivity gradient and those ignoring conductivity, no sinks or sources appeared or disappeared, 3) both CSD approximations revealed: A) a strong sink in layer I representing most likely summed monosynaptic EPSPs of the ventromedial thalamic afferents; B) a strong sink in layer VI, probably representing summed disynaptic EPSPs on dendrites of layer VI pyramidal cells, generated by axons of upper layer pyramidal cells; and C) a sink in layers V/VI representing probably threesynaptic summed EPSPs on dendrites of layer V/VI pyramidal cells.

Relative conductivity measurements were made by using a slightly modified method earlier used by others [8,10,17]. The modification was made in order to avoid damage to the afferent thalamic input fibers: the reference electrode for the application of constant current pulses was placed in the rat’s mouth and not beneath the cortex. The magnitude of conductivity changes with depth in our experiments in the frontal cortex (superficial versus deep layers: 49.9%) was similar to those reported by Holsheimer [11] for the in vitro hippocampus of rats (pyramidal cell layer versus dendritic stratum radiatum: 42%) and those reported by Rappelsberger and his colleagues for the rabbit’s in vivo visual cortex (see fig. 6 [17]). The least conductivity in the visual cortex Rappelsberger et al. found in layer Ш followed by layer IV, layer П and layer V, in this order [17]. In the frontal cortex the conductivity was fairly homogeneous inside the superficial layers I through IV and throughout the deep layers V to VI. Layer IV of the visual cortex (about 15% of the cortical thickness) can be clearly separated from layers II-III (20%) whereas in the frontal cortex layers II through IV (20%) can not be cytoarchitectonically separated from each other [18,20]. Our relative conductivity measurements correspond very well with the known cytoarchitectonical lamination pattern of the frontal cortex showing a 49.9% lower conductivity in the superficial layers with more densely packed and smaller pyramidal cells compared to the less densely distributed and larger pyramidal cells in the deep layers [18,21].

For in vivo current source density analysis and conductivity measurements it is important to avoid mechanical displacement and compression of the cortical tissue by the movement of the recording electrodes. Therefore only microelectrodes with sharpened tips and fine shafts should be used and advanced very slowly. Large changes of the extracellular conductivity were allways seen in those electrode tracks where the distribution of sinks and sources was also shifted in depth, i.e. when the cortical tissue was compressed and displaced by the advancement of the electrode. Assemblies of multiarray electrodes sometimes used in experiments for current source density analysis are more likely to displace cortical tissue resulting in artificial conductivity changes with depth compared to thin microelectrodes with sharpened tips. This could be perhaps one explanation for the stronger effect of the conductivity gradient on current source density in the visual cortex of rabbits described by Rappelsberger and colleagues [17] which used 8- and 16-channel electrode arrays. They claimed the need of conductivity measurements for current density analysis in every experiment, whereas others using moveable single electrodes have denied the significance of conductivity changes with depth [7,11,12,19]. An additional explanation for the higher variability of relative conductivity measurements in experiments using electrode arrays [17] is the much lower spatial sampling rate (150 to 300 μm steps) compared to 100 μm used by Holsheimer [11] and 50 μm steps used in our experiments. A higher variability of relative conductivity estimations with large and sudden changes of the conductivity due to low spatial sampling or mechanical compression and displacement of the cortical tissue results in higher noise levels of CSD estimations using the presumed conductivity changes compared to those ignoring the conductivity.

In our experiments both CSD approximations revealed strong inward membrane currents (sink) with the shortest latency (0.5 ms) near the cortical surface representing most likely summed monosynaptic EPSPs of the ventromedial thalamic afFerents. This is in correspondence with anatomical reports showing the strongest and most dense projection of ventromedial thalamic afFerents to the upper two thirds of layer I of the frontal cortex [1,9]. In most experiments this sink was about 50-100 μm deeper than our depth estimations of the upper two third of layer I would asume. This could be due to a slight displacement of the cortical tissue by the microelectrode entering the cortex. The next strong sink (about 1 ms later) was found in layer VI, probably representing summed disynaptic EPSPs on the soma, proximal or basal dendrites of layer VI pyramidal cells, generated by axons of upper layer pyramidal cells. We know little about the details of the frontal intracortical circuitry and we can only speculate that projections of upper layer pyramidal cells to layer VI pyramidal cells generate this sink. Such connections would make sense for a fast motor output pathway to the striatum and brainstem motor areas via descending projection neurons in layer VI of the frontal (motor) cortex. The third sink (2 to 2.5 ms) was near the boundary between layer V and layer VI and probably represents summed threesynaptic EPSPs on dendrites of layer V/VI pyramidal cells.

In spite of all the methodological problems, which we should always keep in mind, together with sufficient anatomic data the CSD method provides a useful tool for studies of the cortical microcircuitry and cellular mechanisms of the generation of EEG signals by visualization of net membrane current densities as a function of both latency from stimuli and cortical depth.

